# Allelic variation in Class I HLA determines pre-existing memory responses to SARS-CoV-2 that shape the CD8^+^ T cell repertoire upon viral exposure

**DOI:** 10.1101/2021.04.29.441258

**Authors:** Joshua M. Francis, Del Leistritz-Edwards, Augustine Dunn, Christina Tarr, Jesse Lehman, Conor Dempsey, Andrew Hamel, Violeta Rayon, Gang Liu, Yuntong Wang, Marcos Wille, Melissa Durkin, Kane Hadley, Aswathy Sheena, Benjamin Roscoe, Mark Ng, Graham Rockwell, Margaret Manto, Elizabeth Gienger, Joshua Nickerson, MGH COVID-19 Collection and Processing Team, Amir Moarefi, Michael Noble, Thomas Malia, Philip D. Bardwell, William Gordon, Joanna Swain, Mojca Skoberne, Karsten Sauer, Tim Harris, Ananda W. Goldrath, Alex K. Shalek, Anthony J. Coyle, Christophe Benoist, Daniel C. Pregibon

## Abstract

Effective presentation of antigens by HLA class I molecules to CD8^+^ T cells is required for viral elimination and generation of long-term immunological memory. In this study, we applied a single-cell, multi-omic technology to generate the first unified *ex vivo* characterization of the CD8^+^ T cell response to SARS-CoV-2 across 4 major HLA class I alleles. We found that HLA genotype conditions key features of epitope specificity, TCR α/β sequence diversity, and the utilization of pre-existing SARS-CoV-2 reactive memory T cell pools. Single-cell transcriptomics revealed functionally diverse T cell phenotypes of SARS-CoV-2-reactive T cells, associated with both disease stage and epitope specificity. Our results show that HLA variations influence pre-existing immunity to SARS-CoV-2 and shape the immune repertoire upon subsequent viral exposure.

**One-Sentence Summary:** We perform a unified, multi-omic characterization of the CD8^+^ T cell response to SARS-CoV-2, revealing pre-existing immunity conditioned by HLA genotype.

## Introduction

Elicitation of a robust and durable neutralizing antibody response following immunization of large sections of the population with approved SARS-CoV-2 vaccines is limiting viral transmission and decreasing mortality, providing hope that the global threat from the COVID-19 pandemic is diminishing. However, the appearance of new viral variants warrants continued vigilance. A more complete understanding of the underlying cellular mechanisms that regulate host immunity and guarantee long term protection is required. Infection with SARS-CoV-2 leads to an upper respiratory tract infection, which can be benign or even asymptomatic. If not controlled by the immune response, it can evolve into a lethal pneumonia with immunopathology due to excessive amplification of the innate inflammatory response, complicated by several extra-respiratory manifestations (*1*). While humoral responses play an important role in immunological control of infection, the generation of effective cellular immunity and expansion of cytotoxic CD8^+^ memory T cells is also required to eliminate virally infected cells as shown from the earlier SARS-CoV-1 epidemic, even in the absence of seroconversion (*2-7*).

Several recent studies have focused on the discovery of relevant SARS-CoV-2 epitopes in both CD4^+^ and CD8^+^ T cell responses, leveraging *in silico* predictions, stimulation/expansion with peptide pools (*8-18*), and tetramer binding (*19, 20*). Collectively, these studies identified a number of immunodominant epitopes derived across the viral proteome including structural and non-structural proteins (*8-20*). Interestingly, some of these specificities were also detected in uninfected individuals, suggesting potential cross-reactivity from endemic human coronaviruses (HCoV) to which the population is routinely exposed (*21*), though a direct connection to pre-existing memory cells has not been established.

The breadth and nature of the cellular immune response to SARS-CoV-2 infection is driven by diversity in both TCR repertoire and human leukocyte antigen (HLA) genetics. Mammalian cells express up to six different HLA class I alleles that shape antigen presentation in disease, and allelic diversity has been associated with both disease susceptibility and outcome of viral infections (*22, 23*). There are divergent reports regarding HLA polymorphism and COVID-19 incidence and severity, although the major GWAS studies clearly show no dominant effect of the locus (*24-28*). Together with genetic influences on HLA-associated antigen presentation, the clonal selection of T cell receptors (TCRs) that compose an individual’s repertoire contributes to the nature and dynamics of the antiviral response, including cellular cytotoxicity and memory formation. Interestingly, despite a potential TCR diversity of 10^15^ (*29*), several studies have described “public” T cell responses in COVID-19, where complementarity-determining region (CDR) sequences are conserved within and across individuals (*18, 30*). The extent to which TCR diversity, especially in the context of epitope specificity restricted to HLA, contributes to response is not well understood.

Here, we leverage a unique technology to elucidate, at single-cell resolution, the connection between T cell specificity, HLA variation, conserved features of paired α/β TCR repertoires, and cellular phenotype observed in CD8^+^ T cell responses to SARS-CoV-2 infection. We profiled 108,078,030 CD8^+^ T cells *ex vivo* across 76 acute, convalescent, or unexposed individuals, and identified T cell specificity to 648 epitopes presented by four HLA alleles across the SARS-CoV-2 proteome, few of which are implicated by the current variants of concern. Epitope-specific TCR repertoires were surprisingly public in nature, though we found a high degree of pre-existing immunity associated with a clonally diverse response to HLA-B*07:02, which can efficiently present homologous epitopes from SARS-CoV-2 and HCoVs. Transcriptomic analysis and functional validation confirmed a central memory phenotype and TCR cross-reactivity in unexposed individuals with HLA-B*07:02. Our data suggest a strong association between HLA genotype and the CD8^+^ T cell response to SARS-CoV-2, which may have important implications for understanding herd immunity and elements of vaccine design that are likely to confer long-term immunity to protect against SARS-CoV-2 variants and related viral pathogens.

## Results

### Direct ex-vivo detection and decoding of SARS-CoV-2-specific CD8^+^ T cells

We leveraged single-cell RNA-sequencing with DNA-encoded peptide-HLA tetramers to characterize CD8^+^ T cell responses to SARS-CoV-2 across multiple Class I alleles in subjects with varying degrees of disease severity. The technology illustrated in **Fig. 1a** simultaneously determines the specificity of paired α/β TCR sequences for HLA-restricted epitopes and provides transcriptomic phenotype at single-cell resolution. We designed peptide-HLA tetramer libraries to ensure comprehensive coverage of SARS-CoV-2 and related betacoronaviruses across four class I HLA alleles prevalent in North America (A*02:01, B*07:02, A*01:01, and A*24:02, hereafter A*02, B*07, A*01, and A*24). Library inclusion was determined computationally using predicted HLA binding (NetMHC-4.0 (*31*)) of candidate peptides from a set of all possible 9-mers from the SARS-CoV-2 proteome (40% from structural, 60% from non-structural proteins), potentially immunogenic neopeptides from known SARS-CoV-2 variants, and immunogenic epitopes from SARS-CoV-1. A total of 1,355 SARS-CoV-2 related epitopes were included in the libraries in addition to well-characterized epitopes from common endemic viruses (CMV, EBV, and influenza).

**Fig. 1.**
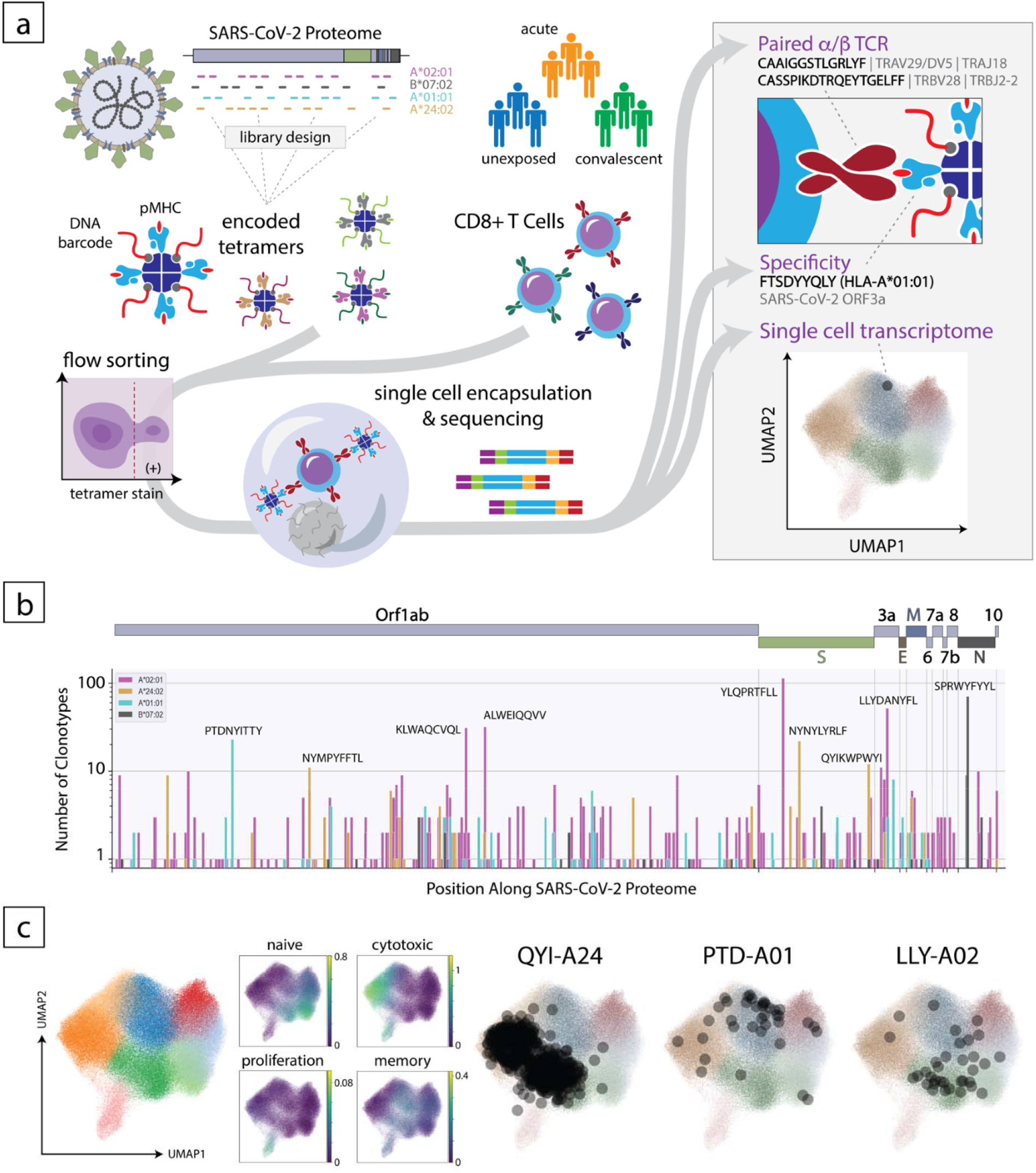
Overview of the experimental approach to decode the CD8^+^ T cell response to SARS-CoV-2. (a) Schematic of the method where encoded tetramer libraries, designed independently for each HLA allele to span the entire SARS-CoV-2-proteome, are used to stain enriched CD8^+^ cells from subject PBMCs, which are then sorted and subjected to single-cell sequencing (left). Using this approach, TCR sequence, peptide/HLA specificity and transcriptomic features are simultaneously acquired for each cell (right). (b) Clonotype specificity detected by HLA allele and epitope across the SARS-CoV-2 proteome. A scheme of the viral ORF structure is shown at the top. Bar colors denote HLA allele. Amino acid sequences of epitopes recognized by the largest number of T cell clonotypes are shown next to the corresponding bar (c). Single-cell transcriptomic analysis showing global UMAP clustering, scoring by functional gene set, and projections onto the transcriptomic UMAP for T cells with specificity toward select epitopes in convalescent individuals. QYI-A24, PTD-A01, and LLY-A02 correspond to QYIKWPWYI in A*24:02, PTDNYITTY in A*01:01, and LLYDANYFL in A*02:01, respectively.

The peptide-HLA tetramer libraries were used to interrogate PBMCs from individuals who had been infected with SARS-CoV-2 (N = 28 convalescent, N = 27 with acute disease that required hospitalization), or who were unexposed (N = 23) as summarized in **Table S1**. For each sample, CD8^+^ cells were isolated from PBMCs (**Methods**), incubated with HLA-matched tetramer libraries, and sorted by flow cytometry to enrich viable, tetramer positive cells. Sorted single cells were encapsulated with DNA-encoded hydrogel beads to provide cell-specific barcodes and unique molecular identifiers (UMIs) that could be used to unify reads across independent sequencing libraries for TCR, peptide-HLA tetramer, and mRNA (**Fig. 1a**). We determined the specificity of TCRs using a classification method that identified UMI counts for TCR-peptide-HLA interactions that were outliers when Z-score transformed within and across cells for each sample (**Methods**). The resulting classifier was evaluated against functional assay data for each allele by a receiver-operator curve (ROC) analysis to identify thresholds, which were then used for normalization. The normalized classifier evaluated by ROC analysis provided an area under the curve (AUC) of 0.82 (**Fig. S1)**, and at a threshold of 1, which was applied to the entire data set, yielded a true positive rate of 93% and a false positive rate of 32%.

From the 55,956,215 CD8^+^ cells interrogated from acute and convalescent COVID-19 patients, we identified high-confidence TCR-peptide-HLA interactions across 434 immunogenic SARS-CoV-2-derived epitopes and 1,163 independent α/β TCR clonotypes (**Fig. 1b, Table S2**). The immunodominant epitopes we discovered *ex vivo* were consistent with those measured by other means (*8-20*), but we also identified many epitopes with less dominant representation (yet observed with two or more reactive clonotypes), 188 of which had not been previously reported as minimal epitopes (**Table S3**). Importantly, CD8^+^ T cell reactivity to SARS-CoV-2 epitopes was observed across the entire proteome, generally distributed in a manner consistent with protein lengths (**Table S4**). Of relevance, 85 of these epitopes were derived from the Spike protein currently used in vaccines, but only six of them (a total of 20 CD8^+^ T cell clonotypes in our study) would be affected by the recent SARS-CoV-2 variants (B.1.1.7, B.1.351, P.1) (**Table S5**).

Dimensionality reduced projections of mRNA expression for 224,780 CD8^+^ T cells revealed the broad phenotypic variance observed within this study spread across 8 clusters (**Fig. 1c**). We defined the phenotypic features of clusters using gene signatures generally associated with various CD8^+^ T cell states, including those with naïve, memory, effector, and proliferative status (**Fig. 1c**). In this space, cells from convalescent patients that recognized different dominant epitopes were commonly associated with divergent phenotypes, as shown for representative epitopes in **Fig. 1c**. For example, T cells specific for QYIKWPWYI in A*24 (QYI-A24) were clustered in regions with high effector scores while those specific for PTDNYITTY in A*01 (PTD-A01) and LLYDANYFL in A*02 (LLY-A02) resided at opposite ends of memory-rich regions. Thus, and as will be further detailed below, the different immunoreactive epitopes of SARS-CoV-2 elicit distinct CD8^+^ T cell phenotypes.

### Evolution of immunoreactivity through COVID-19 disease progression

Having established a broad landscape of SARS-CoV-2-reactive CD8^+^ T cells, we asked how TCR repertoires evolve over the course of infection and recovery. As our approach does not require cell expansion to determine TCR specificity, we were able to directly quantify the frequency of epitope-specific CD8^+^ T cells in the blood of convalescent, acute, and unexposed individuals. **Figure 2a** shows the frequency, for each subject, of T cells reactive to the top five epitopes detected across each of the four HLA variants analyzed. Notably, we observed markedly fewer SARS-CoV-2-specific T cells in patients with acute disease compared to those in convalescence (p = 6.0e-7 for A*02, Wilcoxon rank-sum). The dramatic reduction also applied to memory T cells from prior antiviral responses in these patients, including influenza and EBV, but potentially less to the CMV-specific pool in multiple acute subjects (**Fig. S2**). The paucity of virus-reactive T cells is consistent with the T cell lymphopenia that has been reported to occur in patients with acute COVID-19 (*1, 32*).

**Fig. 2.**
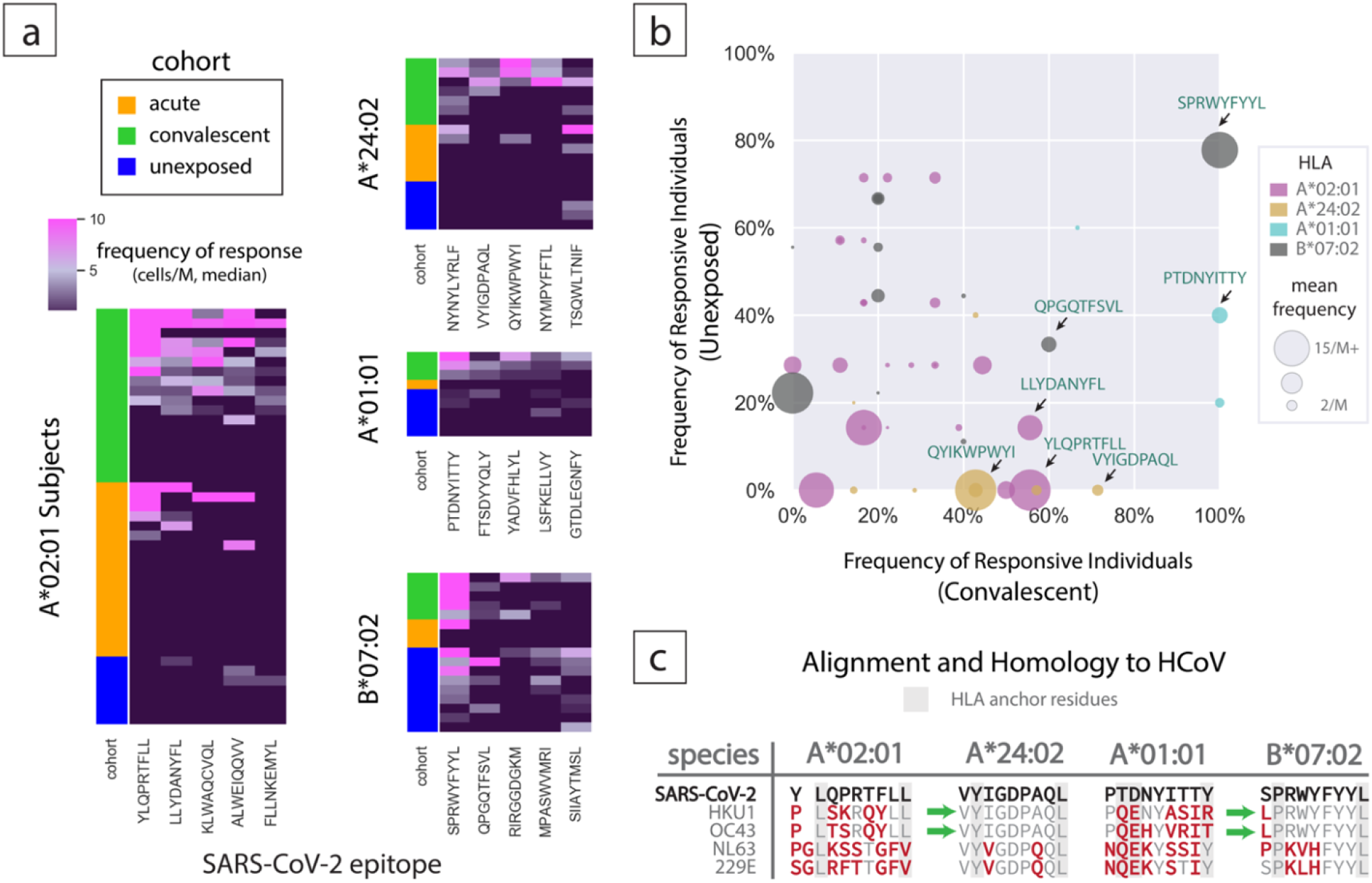
CD8^+^ T cell specificity for major SARS-CoV-2 epitopes across HLA, cohort, and subject. (a) Frequency of T cell response detected (cells per million CD8^+^ cells interrogated) by subject and cohort. Color bar denotes cohort. Heat maps show T cell clonotypes over epitopes and frequency of response as median reactive cells per million T cells for each clonotype (y-axis) and epitope (x-axis) for the indicated HLA alleles. (b) T cell specificity observed in unexposed versus convalescent cohorts represented as percentage of cohort with any detectable frequency of T cell specificity against each epitope. Dots represent epitopes, dot sizes mean combined frequencies across convalescent and unexposed subjects. Sequences of the epitopes with the highest reactivities in convalescent patients are listed. (c) Sequence alignment between select SARS-CoV-2 epitopes and related common cold coronaviruses (HCoV) epitopes. Mismatches are represented in red and HLA anchor residues with a grey background. Green arrows indicate sequences where anchor and all internal residues are conserved between SARS-CoV-2 and HCoV species.

We also observed that the frequencies of SARS-CoV-2-specific T cells in unexposed individuals varied markedly with the HLA allele (**Fig. 2a**). While several dominant epitopes in HLA-A*02, A*24, and A*01 were associated with high-frequency responses in >40% of convalescent subjects (**Fig. 2b**), the depth of the overall response was significantly lower in unexposed compared to convalescent individuals (p=2.3e-5, 2.2e-4, 1.1e-6 by Wilcoxon rank-sum, respectively). In stark contrast, there was no discernible difference in response frequency detected across the most immunodominant epitopes in B*07:02 individuals (p=0.2). In fact, CD8^+^ T cells recognizing nucleocapsid-derived SPRWYFYYL in B*07 (SPR-B07) were found in almost 80% of unexposed subjects with a mean frequency of 4 cells/M cells screened (**Fig. 2b**), presaging the immunodominance of this epitope in convalescent COVID-19 patients, where reactivity was detected in 100% of the samples.

The broad presence of SARS-CoV-2-specific T cells in unexposed B*07 subjects could originate from fortuitous cross-reactivity of a public specificity, or from priming via previous exposure to a highly related endemic human coronavirus (HCoV). Indeed, SPR-B07 shows marked homology to the corresponding segments of the nucleocapsid proteins from multiple prevalent HCoVs, including HKU1 and OC43, with only a single amino acid residue mismatched at the N-terminus (**Fig. 2c**). The nature of the homology preserves internal TCR-contact residues as well as the P and L anchors for HLA binding in peptide positions 2 and 9. Accordingly, the HCoV epitope (LPR-B07) is predicted to bind with high affinity to HLA-B*07 and could reasonably be expected to cross-react with SPR-B07-specific TCRs. Broader sequence alignment with HCoVs revealed very little homology to the immunodominant epitopes of A*02 and A*01 but did identify a perfect match to VYIGDPAQL for A*24 (VYI-A24). Surprisingly, T cell specificity to VYI-A24 was not detected in a single unexposed subject. This likely reflects the lower frequency of response elicited by this epitope or an insufficient commitment to memory following exposure to HCoVs. Overall, we found that the response to SARS-CoV-2 is sharply distinguished by HLA genotype, as can be seen clearly in the case of A*02 and B*07, where it appears that highly specific CD8^+^ responses are either generated *de novo* or amplified from an abundant pre-existing pool, respectively.

### Functional reactivity and cross-reactivity of SARS-CoV2-specific clonotypes

To confirm the specificity and functionality of TCR-peptide-HLA interactions identified in this study, we cloned several of the discovered α/β TCRs clonotypes and expressed them in the TCR-null Jurkat J76 cell line (*33*). Activation of these transductants upon stimulation by SARS-CoV-2 peptides, presented by an HLA-matched lymphoblast cell line, was evaluated by measuring the induction of surface CD69 (**Fig. 3a, Methods**). Altogether, we validated 28 interactions for epitopes derived from Orf1ab, Spike, Nucleocapsid, Membrane, and Orf3a proteins, spanning high confidence interactions observed across multiple cells as well as interactions observed exclusively in single cells (**Table S6**). Dose-response curves for a subset of interactions in A*02 and B*07 are shown in **Figure 3b**. The EC50s measured for these interactions ranged from 1 to 100 nM, with no particular relationship to epitope immunodominance or clonotype frequency measured *ex vivo* from the respective subject. These values are consistent with interactions measured for CMV-specific epitopes in A*02 using the same system. We next used these recombinant TCR expressing cell lines to compare the functional reactivity elicited by homologous epitopes from HCoVs (**Fig. 3c**). Activation was insignificant for the closest homologs of Orf3a-derived LLY-A02 and Orf1ab-derived ALW-A02, all of which actually originated from HCoV spike proteins. In contrast, HKU1 and OC43 homologs of nucleocapsid-derived SPR-B07 and KPR-B07 epitopes drove substantial T cell activation (**Fig. 3c**).

**Fig. 3.**
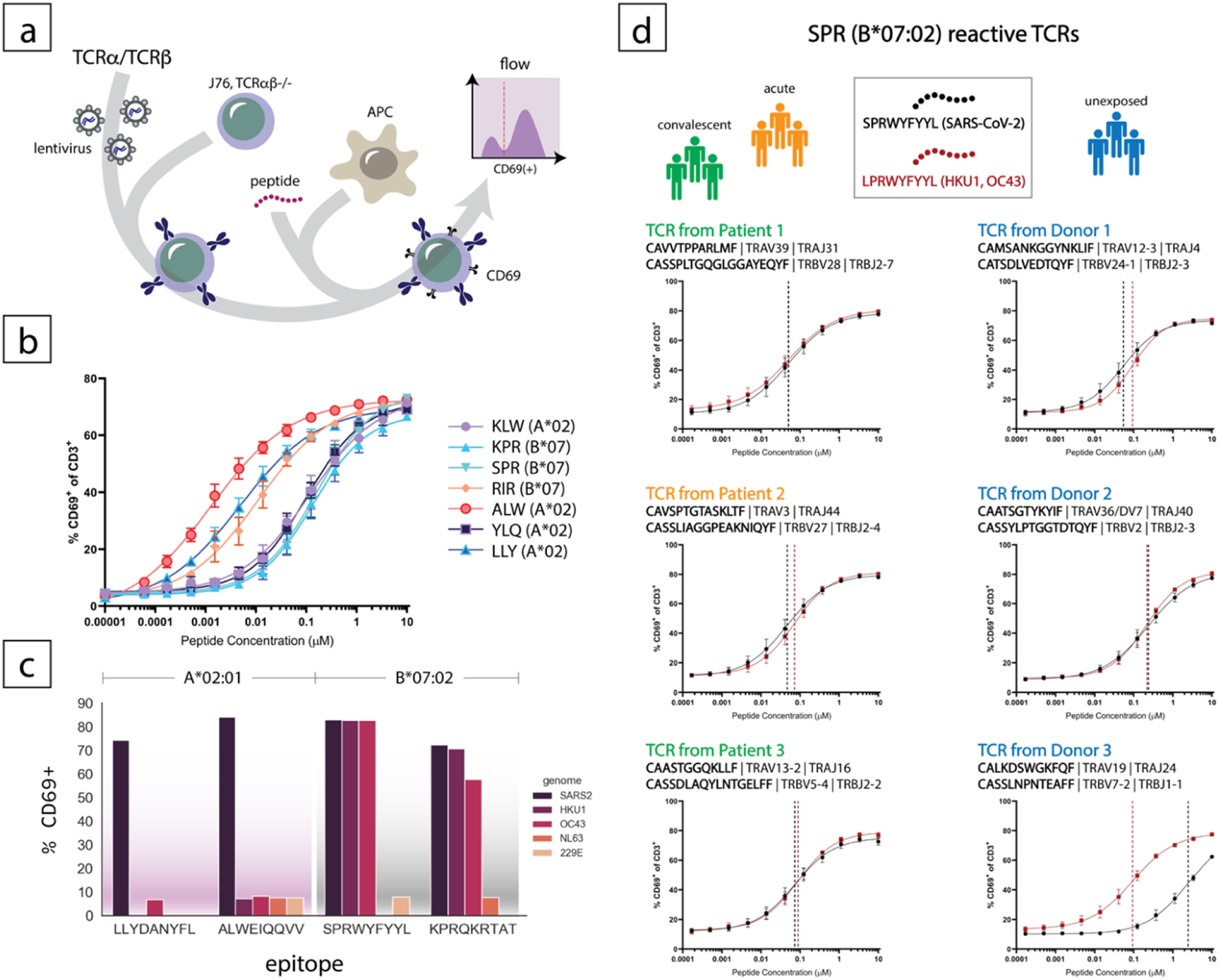
Functional confirmation of identified epitope/HLA interactions with clonotypic TCRs and comparison of SARS-CoV-2 epitopes with common cold coronaviruses. (a) Schematic showing lentiviral transduction of clonotypic recombinant TCRs (rTCRs) into J76 cells, loading of antigen presenting cells (APC) with synthetic peptide, and quantification of activated J76 cells expressing surface CD69. (b) Dose-response curves for TCR-pHLA interactions observed across several canonical epitopes in A*02:01 and B*07:02 backgrounds. Shown are fractions of CD69(+) cells after a 16-hour stimulation. (c) Functional activation of TCRs by canonical and homologous epitopes, represented as fraction of CD69(+) cells after 16-hour stimulation with 10 μM peptide. (d) Dose-response curves for several rTCRs from COVID patients (left) or unexposed subjects (right) stimulated with peptides from SARS-CoV-2 or HCoV HKU1/OC43.

We further assessed the sensitivity of B*07 interactions, comparing the reactivity of SPR-B07-specific clonotypes identified from COVID-19 patients or unexposed subjects to SARS-CoV-2-derived SPR-B07 or HCoV-derived LPR-B07 (**Fig. 3d**). The three TCRs identified from COVID-19 individuals yielded EC50s that were essentially identical for the two epitopes, all falling between 50-100 nM (**Fig. 3d, left**). Two of the TCRs from unexposed individuals yielded EC50s in the same range, again comparable for the HCoV and SARS-CoV-2 variants, while a third showed a >10-fold preference for the HCoV epitope (even though it was originally detected as binding to the SARS-CoV-2 peptide). Aside from providing validation that the specificities detected in our barcoded tetramer technology indeed correspond to antigen-reactive T cells, these findings support that the homologies between SARS-CoV-2 and HCoV epitopes are functionally relevant, and that pre-existing cellular reactivity to SARS-CoV-2 in B*07 subjects likely result from previous exposure to HCoVs like HKU1 or OC43.

### HLA Restricted SARS-CoV-2 Epitopes Impact V(D)J Gene Usage

Given the comprehensive landscape of TCR specificity determined with our approach, we sought to elucidate the extent to which TCR usage is shared within and across subjects. We examined the linkage between paired TCR α/β sequences and their epitope specificity to determine if any features are implicated in the CD8^+^ T cell response to SARS-CoV-2. We used TCRs from 2,469 SARS-CoV-2-specific T cells to perform network mapping of epitope-specific subsets across several immunodominant epitopes identified (**Fig. 4a**). Importantly, because it is known that during development, a TCR β-chain can be paired with many different α-chains, the network analysis allowed clonotype linkages by α or β CDR3 sequences (indicated by edges), identifying conserved motifs based on physicochemical similarity (via BLOSUM matrices) within in the epitope specific T cell population (*34*). T cells from COVID-19 patients that recognize the most dominant A*02-, A*24-, and A*01-restricted epitopes, which have no counterpart in unexposed repertoires, showed a high degree of motif sharing with the exception of KLW-A02 (**Fig. 4a**). Interestingly, all of these epitopes, including KLW-A02, show dominant usage of a single TCR alpha variable (TRAV) region, and in the cases of QYI-A24 and PTD-A01, dominant usage of both TRAV and TCR beta variable (TRBV) regions (**Fig. 4b)**. In marked contrast, SPR-B07-specific T cells, including those that also recognize homologs from HCoV, were far more diverse in CDR3 across subjects (**Fig. 4a**), using 8 TRAV and 3 TRBV regions to cover 50% of the clonotypes represented. We observed two instances of CDR3 homology shared across cohorts, as indicated by the presence of nodes with unconnected edges, which are represented in both network maps.

**Fig. 4.**
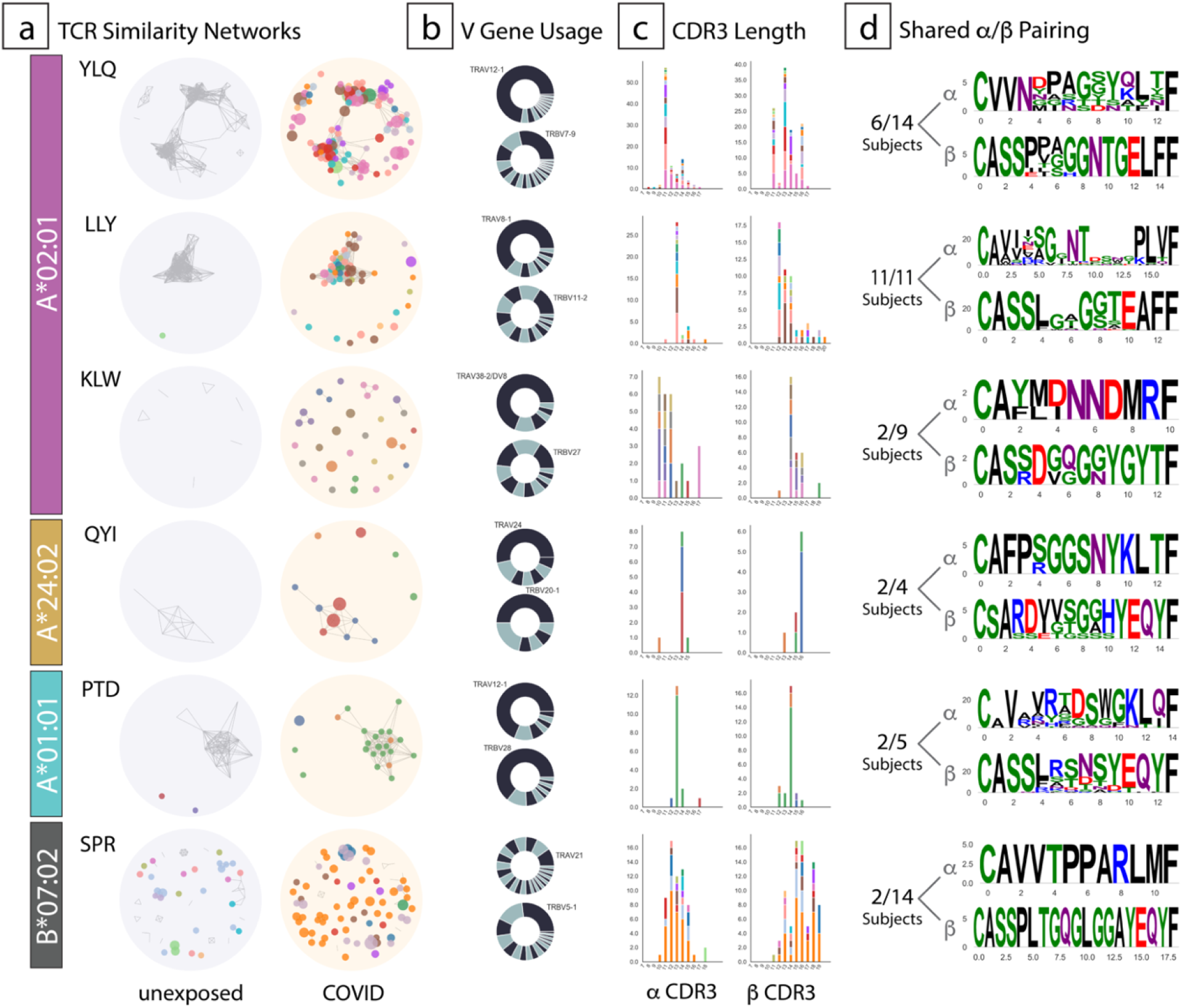
Analysis of TCR sequences from cells specific for the most immunodominant epitopes for each HLA allele tested. (a) Network plots showing biochemical similarity of TCRα or TCRβ CDR3 regions in unexposed subjects (left) or COVID-19 patients (right). Unique subjects are identified by node color. Each node is a unique clonotype within a subject, and the size of the node represents the relative frequency of the response detected. Edges drawn between nodes represent CDR3 homology, and the size of each node represents relative cell frequency. (b) TCRα or TCRβ (V) gene usage across all sequences represented in (a) with the most frequently used gene labeled. (c) Distributions of CDR3 lengths. (d) Paired CDR3 motifs for the most interconnected nodes identified in the network analysis (a).

These comparisons show that the reactivities that appear during SARS-CoV-2 infection may stem from both the amplification of highly related TCRs, or from the usage of diverse pre-existing T cell populations. This conclusion extended to CDR3 lengths (**Fig 4c**), which were tightly distributed for α− and/or β-chains in T cells reactive to the top epitopes in A*02, A*24, and A*01, but significantly less so for SPR-B07. To further elucidate the extent of the public nature of paired α/β TCR usage in COVID-19, we generated consensus sequences from select interconnected network clusters (**Fig. 4d**). This representation provides insight into α/β linkage in the context of public responses that cannot be afforded by bulk sequencing approaches. Most motifs were represented by multiple sequences and shared by 50% of the subjects studied, with the exception of KLW-A02 that was shared across only 22%, and SPR-B07 that was shared across only 14%, notably with identical α/β sequences (**Fig. 4d**). Thus, we have observed divergent TCR repertoire utilization, conditioned by HLA and the presence of diverse, pre-existing reactivity resulting from prior viral exposure.

### CD8^+^ Memory T cell Phenotypes vary with recognition of SARS-CoV-2 epitopes

To examine how CD8^+^ T cell phenotype varied in relation to disease status, HLA/epitope specificity, and TCR diversity, we performed a more detailed analysis of the single-cell transcriptomic data. We leveraged, as an internal reference, the transcriptomic phenotype of T cells reactive to common acute and latent infections, including influenza, EBV, and CMV. To relate these data to existing knowledge on differentially expressed genes that delineate CD8^+^ T subsets, we used supervised partition clustering based on imputed expression (**Methods**) of a set of 51 curated transcripts characteristic of naïve, memory, effector, or chronically-activated/exhausted populations (**Fig. 5a**). This resulted in the identification of seven distinct cell clusters. Some were easily assigned (naïve cells in C1, central memory in C2, and fully activated cytotoxic effectors in C7). Other memory/effector intermediates were more tentatively labeled, as they did not easily fit into existing categorizations (*35-37*). These included a puzzling population (C3, here “CD127^+^ Memory”), which expresses markers of naïve, memory and effector cells, and 3 other clusters with characteristics of memory or chronically activated cells (C4-6).

**Fig. 5.**
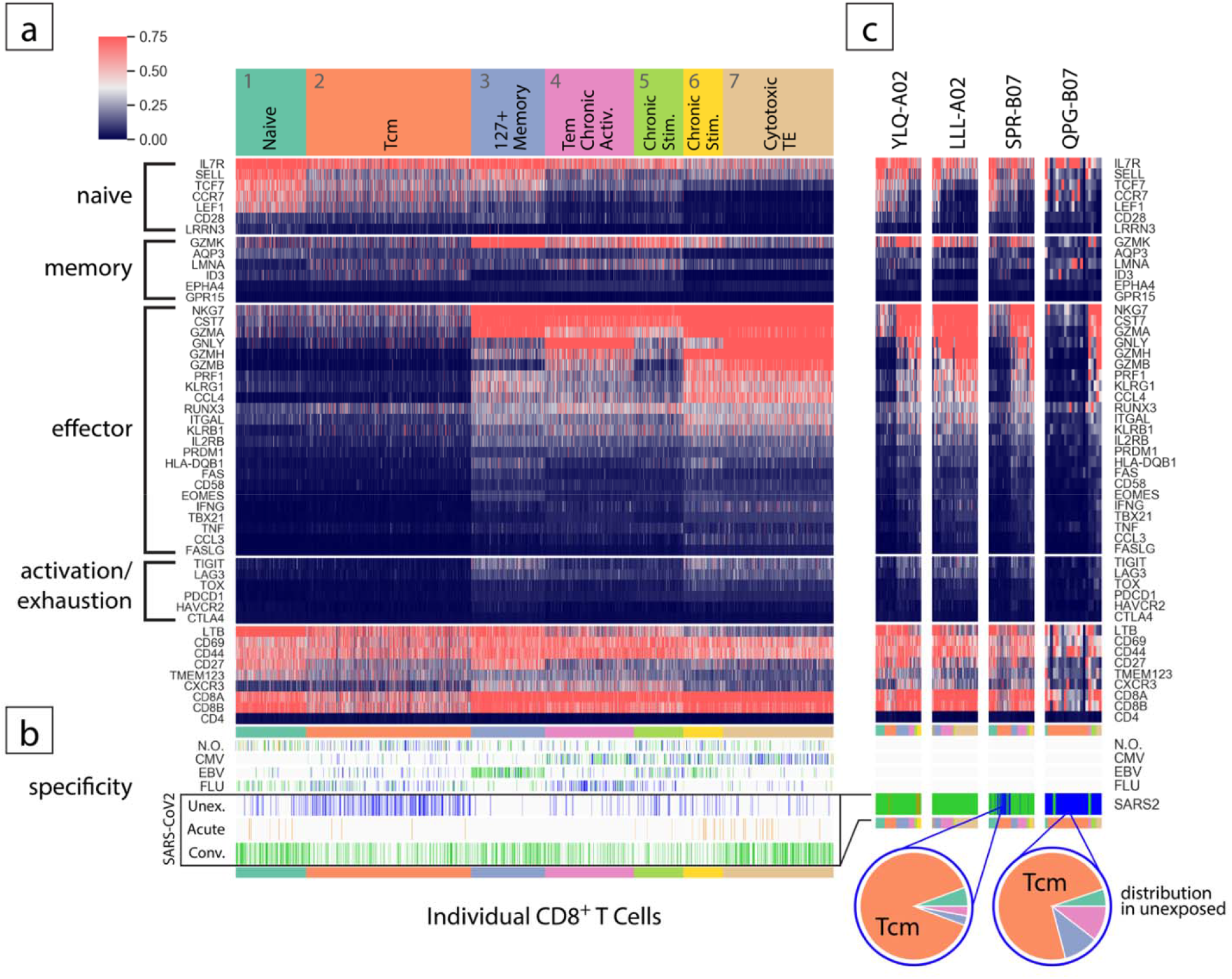
Transcriptomic clustering of T cells based on function-specific gene sets. (a) Single cell gene expression heat map of single CD8^+^ T cells specific for SARS-CoV-2, CMV, EBV, Influenza, or with no observed (N.O.) specificity. Units are ln(TP10K)(*51*). Kmeans clustering identified seven distinct clusters showing gene expression consistent with a range of functional states. (b) Ticks indicate the location and cohort assignment of individual cells with specificity for epitopes from SARS-CoV-2, CMV, EBV, influenza or no observed (N.O.) specificity. (c) Gene expression of single cells with individual epitope specificities indicated in the heat map. In cases where specificity was detected in the unexposed cohort, pie charts are shown to indicate the fraction of cells corresponding to each cluster identified in (a).

SARS-CoV-2 specific T cells were found in all clusters (**Fig. 5b**, bottom), but at proportions that varied with stage of disease and epitope specificity (**Table S7**). Cells from acute patients predominantly showed effector phenotypes, but also paradoxically naïve types. In convalescent donors, T cells from several epitope specificities were broadly distributed, consistent with the resolution of an infection. Several epitope-specific T cell pools were predominantly found in central memory (C2), including PTD-A01 (49%) and LLY-A02 (42%), while others predominantly resided in the cytotoxic terminal effector cluster (C7), including TLM-A02 (80%), and LLL-A02 (61%). In most other reactivities, including SPR-B07, transcriptional profiles in convalescent patients were fairly broadly distributed across all clusters. In contrast, the reactivity in unexposed subjects was dominated by the central memory pool, confirming that the CD8^+^ cells likely result from long-term exposure to cross-reactive antigens. This was especially clear in the case of B*07, where epitope-specific T cells for SPR-B07, QPG-B07, and SII-B07 were represented in central memory (C2) at proportions of 88%, 75%, and 67%, respectively. Other notable reactivities associated with central memory include TSQ-A24 (70%) and NSS-A01 (68%), though the source of these memory cells, like QPG-B07 and SII-B07, does not appear to be from HCoV exposure based on a lack of homology. Overall, this analysis provides further evidence that SPR-B07 responses to SARS-CoV-2 are likely drawn from a pre-existing memory pool and that commitment to different cell fate is dependent on epitope specificity.

We also observed some interesting dynamics between SARS-CoV-2 infection and existing T cell pools specific for common viral infections, with differentiated outcomes likely shaped by exposure history (**Fig. 5b**). Influenza-specific CD8^+^ T cells, which result from vaccination or past infections, mapped primarily to the central memory (C1) and effector memory (C3) compartments in unexposed individuals. Proportions were stable across epitope specificities in COVID-19 patients with the exception of GIL-A02, where the proportion of effector memory cells decreased from 50% to 0% and a naïve population representing 30% of the cells paradoxically emerged. CMV-and EBV-specific T cells, likely subject to more chronic stimulation from low-level re-activation of these integrated herpesviruses, mapped to more activated pools in unexposed subjects, as has been described by others (*38*). After SARS-CoV-2 infection, EBV-specific cells shifted markedly from central memory (C2) and chronically stimulated compartments (C5) into the 127^+^ memory cluster (C3). These changes may reflect either bystander activation, perhaps as a result of the high cytokine release in COVID-19 patients, or from changes in homing or recirculation patterns that bring into the blood cells normally sequestered in tissues. These observations suggest that, in addition to inducing lymphopenia, COVID-19 strongly reshuffles third-party antiviral T cell pools, the extent of which may be associated with exposure history and, at least to some degree, epitope specificity.

## Discussion

Here we presented the first unified description of the CD8^+^ T cell response to SARS-CoV-2, highlighting the importance of HLA genetics, TCR repertoire diversity, and epitope-specific navigation through a complex transcriptomic phenotype at various stages of disease. In building a comprehensive map of immunodominant, HLA-restricted epitopes broadly derived from proteins across the entire SARS-CoV-2 proteome, we highlight how only some HLA haplotypes are associated with the existence of a pre-existing CD8^+^ T cell memory pool in unexposed individuals. We further show how HLA variation plays an important role in shaping the diversity of CD8^+^ T cell repertoires upon exposure to SARS-CoV-2, and that cellular phenotype and commitment to memory can be associated with epitope-specificity in the context of both SARS-CoV-2 and latent EBV infections.

The presence of SARS-CoV-2 reactive CD8^+^ T cells has been linked to milder disease (*5, 11, 12*), although the precise link between cellular immunity and host protection still remains to be further understood (*7, 39, 40*). We found that individuals carrying HLA-B*07 show a CD8^+^ T cell response that is dominated by pre-existing memory pools reactive to multiple SARS-CoV-2 epitopes, especially SPR-B07, which is likely induced by previous exposures to benign HCoVs. In contrast, the immunodominant responses in A*02 individuals (e.g. to YLQ-A02, LLY-A02) are driven largely by the expansion of antigen-inexperienced SARS-CoV-2-specific T cells. It is interesting to note that CD8^+^ T cell cross-reactivity may be less widespread in unexposed individuals than for CD4^+^ T cross-reactivity, for which ∼50% of unexposed individuals exhibited CD4^+^ T cell memory (*16*). Our data provides a basis for this limited representation of the CD8^+^ T cell repertoire in that only a subpopulation of individuals carrying a specific HLA allele would have these pre-existing memory CD8^+^ T cells.

The interplay between HLA-restricted epitope presentation and available TCR repertoire shapes the cellular response to SARS-CoV-2. There are few limited studies suggesting an influence of HLA genotype on COVID-19 severity (*27, 41-43*). Large-scale, high-resolution HLA mapping, consistent with what we have done for select HLAs in this work, may help identify relationships between HLA genotype and protection against severe disease, ideally uncovering mechanism. Here, we observed an interesting connection between TCR repertoire diversity and HLA restriction. Responses seen in A*02, A*24, and A*01 were more often associated with “public” CDR3 motifs and consistent V gene segment usage in the α− and/or β− chains. In contrast, the dominant immune response in B*07 leveraged a significantly more diverse TCR repertoire. Several contributors to public TCR responses have been proposed, focusing on the physicochemical features of HLA-restricted peptides (e.g. “featureless” peptide-HLAs may drive a public response) and convergent recombination of TCR dimers (*44*). The method described in this work provides an ideal system to address this question. Perhaps counterintuitively, our results show that in the case of COVID-19, the largest pool of potentially protective, pre-existing cellular immunity is derived from one of the *least* public epitope-specific repertoires, possibly reflecting the influence of repeated acute infections with HCoVs throughout the life of the individuals.

Beyond the comprehensive deciphering of TCR specificity reported here, we also provided a detailed picture of the complex and dynamic transcriptional landscape of the CD8^+^ T response to SARS-CoV-2. Importantly, we were able to demonstrate that the pre-existing SPR-B07 reactivity, observed in ∼80% of unexposed subjects with HLA-B*07, was predominantly associated with a central memory-like transcriptional profile (88%), confirming that it originates from prior exposures. In convalescent patients, we observed a much broader distribution of SPR-B07-reactive T cells spanning every functional state at proportions ranging from 5-29% (**Table S7**). This is consistent with late contraction/early memory formation described for SARS-CoV-2 in a recent study (*12*), where cells spanned naïve, central memory, various classifications of effector memory, and terminally differentiated effector memory expressing RA (TEMRA). There was no evidence for a particularly frequent “exhausted” state among SARS-CoV-2-specific CD8^+^ T cells, as suggested elsewhere (*45, 46*) (acknowledging that the phenotypic state is a proxy for true reactivity testing, and that blood T cells may not fully reflect what happens in the lung). We also did not find evidence of “antigenic sin” resulting from HCoV pre-exposure (*47*) that would stifle an effective response to SARS-CoV-2-unexposed B*07 individuals. It will be interesting to determine whether HLA haplotype plays a role in the durability of the CD8^+^ T cell responses, especially to SARS-CoV-2 vaccines, which may have profound impact for long term protection across different ethnic groups and geographic regions.

Another interesting observation from this work, as noted by others (*48*), is that even at the height of infection or shortly after viral clearance, the cumulative anti-SARS-CoV-2 CD8^+^ T cell response barely reached the frequency of anti-influenza memory responses and was well below the frequencies that could be achieved by CMV-specific cells in the same individuals (**Fig. S2**). This was particularly evident in the acutely infected individuals, at a time where the contribution of cytotoxic CD8^+^ T cells would have been most important. We acknowledge the caveat that peripheral frequencies were measured, and some degree of sequestration in viral target tissues, such as the lung, is likely to occur in acute patients. Yet, the response seems much more muted than the “all hands-on deck” observed in some other viral infections (*49*). This meager outcome was seen both for the cross-reactive “secondary responses” by memory T cells pre-primed by endemic HCoVs, as well as for the primary responses of truly SARS-CoV-2 species-specific CD8^+^ T cells amplified *de novo*. This suggests that the paucity likely does not result from a blocking of primary activation, but from a dampening of all specific CD8^+^ T cells. Consistent with this notion, frequencies of influenza/EBV/CMV reactive cells were also lower in acute COVID-19 patients, compared to SARS-CoV-2 “naïve” individuals. It has been proposed that the lethal cytokine storm in severe COVID-19 stems from innate immune functions overcompensating for adaptive immune system failures (*2*). In this line of reasoning, one might propose that SARS-CoV-2’s noxiousness stems from a broad obstruction of antiviral CD8^+^ T cell responses. We have recently described a “super-Treg” phenotype in severe COVID-19, with heightened expression of FoxP3 and Treg effector molecules, akin to tumor-infiltrating Treg cells (*50*), and one possible interpretation is that overactive Tregs are overly suppressing these CD8^+^ T cells in severe COVID-19 patients.

Given the widespread lymphopenia observed in acute COVID-19, we pondered the possibility of latent virus reactivation with the loss of protective CMV-and EBV-specific T memory pools. While we have no direct evidence of impact on disease outcome, we do see a significant alteration of cell state within these subsets. While CMV-reactive cells remained within, though somewhat shuffled, the same effector/memory transcriptional phenotypes between unexposed and COVID-19 cohorts (including chronic stimulation, cytotoxic terminal effector, and terminal effector memory), we observed a dramatic shift of EBV-specific cells from chronic stimulation and central memory into the interesting “127^+^ memory” state in COVID-19-exposed individuals. These cells expressed moderate to high levels of many naïve (IL7R, SELL, CCR7), memory (GZMK), and effector-associated genes (NKG7, CST7, GZMA), along with markers of activation/exhaustion (TIGIT, LAG3), making them particularly interesting and difficult to ascribe to conventional phenotype labels. Recently, two new transcriptionally distinct stem-like CD8^+^ T cell memory states were described, one of which was functionally committed to a dysfunctional lineage (*37*). As these cell states were differentiated by many of the same markers observed in our “127^+^ memory” compartment, it would be interesting to see to what extent these “127^+^ memory” cells, dominated by EBV-reactive pools, experience similar fates of dysfunction. We speculate that this phenotype may be a consequence of the particular inflammatory milieu of COVID-19 patients.

In conclusion, we leveraged a powerful single-cell technology to better elucidate the roles of HLA variation, TCR diversity, and cellular phenotypes in establishing pre-existing immunity to SARS-CoV-2. We observed the presence of a diverse and immuno-dominant nucleocapsid epitope-specific memory pool in subjects with HLA-B*07 but saw little evidence of similar reactivity in individuals with other HLA alleles. Outside of the HLA-B*07, the epitope-specific TCR repertoires observed were largely public in nature. We measured a diverse landscape of T cell phenotypes associated with SARS-CoV-2 infection, and also observed an influence on T cell repertoires reactive to persistent and latent infections with other viruses. Overall, this work provides a framework for the unified characterization of the cellular response to novel viral infections. The ability to understand the basis of cellular immunity to SARS-CoV-2 and other pathogens will provide insight for the continued assessment of immune surveillance, health security, and long-term protection from future respiratory pathogens.

## Author Contributions

Experimental design, J.M.F., D.L-E, A.D., G.L., V.R, J.L., C.D., V.R., A.M., M.N., K.S., T.H., A.C., C.B., D.C.P. Reagents and samples, D.L-E, C.T., J.L., C.D., A.H., V.R., Y.W., M.W., M.D., B.R., M.N., M.M., E.G., J.N., MGH COVID-19 Collection and Processing Team, T.M., P.B., W.G., J.S. Analysis, J.M.F, D.L-E., A.D., G.L., J.L, C.D.,K.H. A.S., G.R., M.N., M.S., K.S., T.H., A.W.G., A.K.S., A.C., C.B., D.C.P. Writing, J.M.F., A.D., A.K.S., A.C., C.B., D.C.P.

## Data and materials availability

All data are available in the manuscript or supplementary materials. All reasonable request for data and material will be fulfilled.

## MGH COVID-19 Collection and Processing Team participants

Nikolaus Jilg^1^, Jonathan Li^1^, Alex Rosenthal^1^, Colline Wong^1^, George Daley^2^, David Golan^2^, Howard Heller^2^, Arlene Sharpe^2^, Betelihem A. Abayneh^3^, Patrick Allen^3^, Diane Antille^3^, Katrina Armstrong^3^, Siobhan Boyce^3^, Joan Braley^3^, Karen Branch^3^, Katherine Broderick^3^, Julia Carney^3^, Andrew Chan^3^, Susan Davidson^3^, Michael Dougan^3^, David Drew^3^, Ashley Elliman^3^, Keith Flaherty^3^, Jeanne Flannery^3^, Pamela Forde^3^, Elise Gettings^3^, Amanda Griffin^3^, Sheila Grimmel^3^, Kathleen Grinke^3^, Kathryn Hall^3^, Meg Healy^3^, Deborah Henault^3^, Grace Holland^3^, Chantal Kayitesi^3^, Vlasta LaValle^3^, Yuting Lu^3^, Sarah Luthern^3^, Jordan Marchewka Schneider^3^, Brittani Martino^3^, Roseann McNamara^3^, Christian Nambu^3^, Susan Nelson^3^, Marjorie Noone^3^, Christine Ommerborn^3^, Lois Chris Pacheco^3^, Nicole Phan^3^, Falisha A Porto^3^, Edward Ryan^3^, Kathleen Selleck^3^, Sue Slaughenhaupt^3^, Kimberly Smith Sheppard^3^, Elizabeth Suschana^3^, Vivine Wilson^3^, Mary Carrington^4^, Maureen Martin^4^, Yuko Yuki^4^, Galit Alter^5^, Alejandro Balazs^5^, Julia Bals^5^, Max Barbash^5^, Yannic Bartsch^5^, Julie Boucau^5^, Mary Carrington^5^, Josh Chevalier^5^, Fatema Chowdhury^5^, Edward DeMers^5^, Kevin Einkauf^5^, Jon Fallon^5^, Liz Fedirko^5^, Kelsey Finn^5^, Pilar Garcia-Broncano^5^, Musie S. Ghebremichael^5^, Ciputra Hartana^5^, Chenyang Jiang^5^, Kelly Judge^5^, Paulina Kaplonek^5^, Marshall Karpell^5^, Peggy Lai^5^, Evan C. Lam^5^, Kristina Lefteri^5^, Xiaodong Lian^5^, Mathias Lichterfeld^5^, Daniel Lingwood^5^, Hang Liu^5^, Jinqing Liu^5^, Natasha Ly^5^, Zachary Manickas Hill^5^, Ashlin Michell^5^, Ilan Millstrom^5^, Noah Miranda^5^, Claire O’Callaghan^5^, Matthew Osborn^5^, Shiv Pillai^5^, Yelizaveta Rassadkina^5^, Alexandra Reissis^5^, Francis Ruzicka^5^, Kyra Seiger^5^, Libera Sessa^5^, Christianne Sharr^5^, Sally Shin^5^, Nishant Singh^5^, Weiwei Sun^5^, Xiaoming Sun^5^, Hannah Ticheli^5^, Alicja Trocha-Piechocka^5^, Bruce Walker^5^, Daniel Worrall^5^, Xu

G. Yu^5^, Alex Zhu^5^

^1^ Brigham and Women’s Hospital, Boston, MA, USA

^2^ Harvard Medical School, Boston, MA, USA

^3^ Massachusetts General Hospital, Boston, MA, USA

^4^ National Cancer Institute, Frederick, MD, USA

^5^ Ragon Institute of MGH, MIT and Harvard, Cambridge, MA, USA

## SUPPLEMENTARY MATERIALS

Materials and Methods Figs. S1 to S2

Tables S1 to S7

## REFERENCES AND NOTES

1. C. Huang et al., Clinical features of patients infected with 2019 novel coronavirus in Wuhan, China. Lancet 395, 497–506 (2020).

2. A. Sette, S. Crotty, Adaptive immunity to SARS-CoV-2 and COVID-19. Cell 184, 861–880 (2021).

3. O. W. Ng et al., Memory T cell responses targeting the SARS coronavirus persist up to 11 years post-infection. Vaccine 34, 2008–2014 (2016).

4. J. Seow et al., Longitudinal observation and decline of neutralizing antibody responses in the three months following SARS-CoV-2 infection in humans. Nat Microbiol 5, 1598–1607 (2020).

5. C. Rydyznski Moderbacher et al., Antigen-Specific Adaptive Immunity to SARS-CoV-2 in Acute COVID-19 and Associations with Age and Disease Severity. Cell 183, 996–1012 e1019 (2020).

6. J. M. Dan et al., Immunological memory to SARS-CoV-2 assessed for up to eight months after infection. bioRxiv, (2020).

7. K. Sauer, T. Harris, An Effective COVID-19 Vaccine Needs to Engage T Cells. Front Immunol 11, 581807 (2020).

8. Y. Peng et al., Broad and strong memory CD4(+) and CD8(+) T cells induced by SARS-CoV-2 in UK convalescent individuals following COVID-19. Nat Immunol 21, 1336–1345 (2020).

9. J. Braun et al., SARS-CoV-2-reactive T cells in healthy donors and patients with COVID-19. Nature 587, 270–274 (2020).

10. A. Nelde et al., SARS-CoV-2-derived peptides define heterologous and COVID-19-induced T cell recognition. Nat Immunol 22, 74–85 (2021).

11. T. Sekine et al., Robust T Cell Immunity in Convalescent Individuals with Asymptomatic or Mild COVID-19. Cell 183, 158–168 e114 (2020).

12. I. Schulien et al., Characterization of pre-existing and induced SARS-CoV-2-specific CD8(+) T cells. Nat Med 27, 78–85 (2021).

13. J. Mateus et al., Selective and cross-reactive SARS-CoV-2 T cell epitopes in unexposed humans. Science 370, 89–94 (2020).

14. A. P. Ferretti et al., Unbiased Screens Show CD8(+) T Cells of COVID-19 Patients Recognize Shared Epitopes in SARS-CoV-2 that Largely Reside outside the Spike Protein. Immunity 53, 1095–1107 e1093 (2020).

15. A. Tarke et al., Comprehensive analysis of T cell immunodominance and immunoprevalence of SARS-CoV-2 epitopes in COVID-19 cases. bioRxiv, (2020).

16. A. Grifoni et al., Targets of T Cell Responses to SARS-CoV-2 Coronavirus in Humans with COVID-19 Disease and Unexposed Individuals. Cell 181, 1489–1501 e1415 (2020).

17. D. Weiskopf et al., Phenotype and kinetics of SARS-CoV-2-specific T cells in COVID-19 patients with acute respiratory distress syndrome. Sci Immunol 5, (2020).

18. T. M. Snyder et al., Magnitude and Dynamics of the T-Cell Response to SARS-CoV-2 Infection at Both Individual and Population Levels. medRxiv, (2020).

19. H. Kared et al., CD8+ T cell responses in convalescent COVID-19 individuals target epitopes from the entire SARS-CoV-2 proteome and show kinetics of early differentiation. bioRxiv, (2020).

20. S. K. Saini et al., SARS-CoV-2 genome-wide T cell epitope mapping reveals immunodominance and substantial CD8(+) T cell activation in COVID-19 patients. Sci Immunol 6, (2021).

21. G. J. Gorse, G. B. Patel, J. N. Vitale, T. Z. O’Connor, Prevalence of antibodies to four human coronaviruses is lower in nasal secretions than in serum. Clin Vaccine Immunol 17, 1875–1880 (2010).

22. K. S. MacDonald et al., Influence of HLA supertypes on susceptibility and resistance to human immunodeficiency virus type 1 infection. J Infect Dis 181, 1581–1589 (2000).

23. E. E. Ochoa et al., HLA-associated protection of lymphocytes during influenza virus infection. Virol J 17, 128 (2020).

24. G. G. Severe Covid et al., Genomewide Association Study of Severe Covid-19 with Respiratory Failure. N Engl J Med 383, 1522–1534 (2020).

25. E. Pairo-Castineira et al., Genetic mechanisms of critical illness in COVID-19. Nature 591, 92–98 (2021).

26. J. R. Habel et al., Suboptimal SARS-CoV-2-specific CD8(+) T cell response associated with the prominent HLA-A*02:01 phenotype. Proc Natl Acad Sci U S A 117, 24384–24391 (2020).

27. M. Shkurnikov et al., Association of HLA Class I Genotypes With Severity of Coronavirus Disease-19. Front Immunol 12, 641900 (2021).

28. A. Nguyen et al., Human Leukocyte Antigen Susceptibility Map for Severe Acute Respiratory Syndrome Coronavirus 2. J Virol 94, (2020).

29. Q. Qi et al., Diversity and clonal selection in the human T-cell repertoire. Proc Natl Acad Sci U S A 111, 13139–13144 (2014).

30. A. S. Shomuradova et al., SARS-CoV-2 Epitopes Are Recognized by a Public and Diverse Repertoire of Human T Cell Receptors. Immunity 53, 1245–1257 e1245 (2020).

31. M. Andreatta, M. Nielsen, Gapped sequence alignment using artificial neural networks: application to the MHC class I system. Bioinformatics 32, 511–517 (2016).

32. G. Chen et al., Clinical and immunological features of severe and moderate coronavirus disease 2019. J Clin Invest 130, 2620–2629 (2020).

33. S. Jutz et al., Assessment of costimulation and coinhibition in a triple parameter T cell reporter line: Simultaneous measurement of NF-kappaB, NFAT and AP-1. J Immunol Methods 430, 10–20 (2016).

34. P. Dash et al., Quantifiable predictive features define epitope-specific T cell receptor repertoires. Nature 547, 89–93 (2017).

35. P. A. Szabo et al., Single-cell transcriptomics of human T cells reveals tissue and activation signatures in health and disease. Nat Commun 10, 4706 (2019).

36. G. Monaco et al., RNA-Seq Signatures Normalized by mRNA Abundance Allow Absolute Deconvolution of Human Immune Cell Types. Cell Rep 26, 1627–1640 e1627 (2019).

37. G. Galletti et al., Two subsets of stem-like CD8(+) memory T cell progenitors with distinct fate commitments in humans. Nat Immunol 21, 1552–1562 (2020).

38. S. P. H. van den Berg et al., The hallmarks of CMV-specific CD8 T-cell differentiation. Med Microbiol Immunol 208, 365–373 (2019).

39. A. Addetia et al., Neutralizing Antibodies Correlate with Protection from SARS-CoV-2 in Humans during a Fishery Vessel Outbreak with a High Attack Rate. J Clin Microbiol 58, (2020).

40. A. Fontanet, S. Cauchemez, COVID-19 herd immunity: where are we? Nat Rev Immunol 20, 583–584 (2020).

41. C. Liang et al., Population-Predicted MHC Class II Epitope Presentation of SARS-CoV-2 Structural Proteins Correlates to the Case Fatality Rates of COVID-19 in Different Countries. Int J Mol Sci 22, (2021).

42. A. Anzurez et al., Association of Human Leukocyte Antigen DRB1*09:01with severe COVID-19. HLA, (2021).

43. P. Correale et al., HLA-B*44 and C*01 Prevalence Correlates with Covid19 Spreading across Italy. Int J Mol Sci 21, (2020).

44. V. Venturi et al., A mechanism for TCR sharing between T cell subsets and individuals revealed by pyrosequencing. J Immunol 186, 4285–4294 (2011).

45. H. Y. Zheng et al., Elevated exhaustion levels and reduced functional diversity of T cells in peripheral blood may predict severe progression in COVID-19 patients. Cell Mol Immunol 17, 541–543 (2020).

46. B. Diao et al., Reduction and Functional Exhaustion of T Cells in Patients With Coronavirus Disease 2019 (COVID-19). Front Immunol 11, 827 (2020).

47. E. L. Brown, H. T. Essigmann, Original Antigenic Sin: the Downside of Immunological Memory and Implications for COVID-19. mSphere 6, (2021).

48. J. C. Law et al., Systematic Examination of Antigen-Specific Recall T Cell Responses to SARS-CoV-2 versus Influenza Virus Reveals a Distinct Inflammatory Profile. J Immunol 206, 37–50 (2021).

49. Z. M. Ndhlovu et al., Augmentation of HIV-specific T cell function by immediate treatment of hyperacute HIV-1 infection. Sci Transl Med 11, (2019).

50. S. Galvan-Pena et al., Profound Treg perturbations correlate with COVID-19 severity. bioRxiv, (2020).

51. T. Stuart et al., Comprehensive Integration of Single-Cell Data. Cell 177, 1888–1902 e1821 (2019).

52. H. L. Oh et al., Engineering T cells specific for a dominant severe acute respiratory syndrome coronavirus CD8 T cell epitope. J Virol 85, 10464–10471 (2011).

53. K. M. Campbell, G. Steiner, D. K. Wells, A. Ribas, A. Kalbasi, Prediction of SARS-CoV-2 epitopes across 9360 HLA class I alleles. bioRxiv, (2020).

54. A. Grifoni et al., A Sequence Homology and Bioinformatic Approach Can Predict Candidate Targets for Immune Responses to SARS-CoV-2. Cell Host Microbe 27, 671–680 e672 (2020).

55. J. D. Altman, M. M. Davis, MHC-Peptide Tetramers to Visualize Antigen-Specific T Cells. Curr Protoc Immunol 115, 17 13 11–17 13 44 (2016).

56. F. A. Wolf, P. Angerer, F. J. Theis, SCANPY: large-scale single-cell gene expression data analysis. Genome Biol 19, 15 (2018).

57. B. Hie, B. Bryson, B. Berger, Efficient integration of heterogeneous single-cell transcriptomes using Scanorama. Nat Biotechnol 37, 685–691 (2019).

58. D. Aran et al., Reference-based analysis of lung single-cell sequencing reveals a transitional profibrotic macrophage. Nat Immunol 20, 163–172 (2019).

59. D. van Dijk et al., Recovering Gene Interactions from Single-Cell Data Using Data Diffusion. Cell 174, 716–729 e727 (2018).

