## Supplementary Methods, Figures and Tables for "Allelic variation in Class I HLA determines pre-existing memory responses to SARS-CoV-2 that shape the CD8^+^ T cell repertoire upon viral exposure"

**This PDF file includes:**

Materials and Methods

Figs. S1 to S2

Tables S4 to S6

Captions for Tables S1 to S3 and S7

**Other Supplementary Materials for this manuscript include the following:**

Tables S1, S2, S3 and S7 (.xlsx)

Materials and Methods

**Antigen library design.** Antigenic peptide libraries were made by scoring all possible 9mer peptides derived from the entire SARS-CoV2 (NC_045512.2) proteome using netMHC-4.0 (*31*) in the HLA-A*02:01, HLA-A*01:01, HLA-A*24:02 or HLA-B*07:02 alleles. SARS-CoV-1 peptides that had evidence of T cell positive assays, obtained from the Immune Epitope Database ([www.iedb.org](http://www.iedb.org); (*52*)), and that were highly homologous to their SARS-CoV2 counterparts within hamming-distance of 2 were converted to 9-mers. Additionally, SARS-CoV2 peptides predicted to raise immunogenic responses by others were also included (*53, 54*). Finally, libraries included a set of well-defined viral epitopes from Cytomegalovirus, Epstein-Barr virus, and Influenza viruses (CEF peptide pool) that elicit T cell responses in the population at large. Antigenic peptides with 500 nM affinity or lower were then selected.

**Production of tetramer library pools.** HLA-A*01:01, -A*02:01, -A*24:02 and HLA-B*07:02 extracellular domains were expressed in *E. coli* and refolded along with beta-2-microglobulin and UV-labile place-holder peptides STAPGJLEY, KILGFVFJV, VYGJVRACL and AARGJTLAM, respectively (*55*). The MHC monomer was then purified by size exclusion chromatography (SEC). MHC tetramers were produced by mixing alkylated MHC monomers and azidylated streptavidin in 0.5 mM copper sulfate, 2.5 mM BTTAA and 5 mM ascorbic acid for up to 4 h on ice, followed by purification of highly multimeric fractions by SEC. Individual peptide exchange reactions containing 500 nM MHC tetramer and 60 μM peptide were exposed to long-wave UV (366 nm) at a distance of 2-5 cm for 30 min at 4˚C, followed by 30 min incubation at 30˚C. A biotinylated oligonucleotide barcode (Integrated DNA Technologies) was added to each individual reaction followed by 30 minute incubation at 4˚C. Individual tetramer reactions were then pooled and concentrated using 30 kDa molecular weight cut-off centrifugal filter units (Amicon).

**Cell Staining.** Peripheral blood mononuclear cells (PBMCs) from convalescent COVID-19 positive donors or unexposed donors were obtained from Precision 4 Medicine (USA), the Massachusetts Consortium on Pathogen Readiness (*MassCPR*), or CTL (USA), all under appropriate informed consent. PBMCs were thawed, and CD8+ T cells were enriched by magnetic-activated cell sorting (MACS) using a CD8+ T Cell Isolation Kit (Miltenyi) following the manufacturers protocol. The CD8+ T cells were then stained with 1 nM final concentration tetramer library in the presence of 2 mg/mL *salmon sperm* DNA in PBS with 0.5% BSA solution for 20 minutes. Cells were then labeled with anti-TCR ADT (IP26, Biolegend) for 15 minutes followed by washing. Tetramer bound cells were then labeled with PE conjugated anti-DKDDDDK-Flag antibody (BioLegend) followed by dead cell discrimination using 7-amino-actinomycin D (7-AAD). The live, tetramer positive cells were sorted using a Sony MA900 Sorter (Sony).

**Single-cell Sequencing.** Tetramer positive cells were counted by Nexcelom Cellometer (Lawrence, MA, USA) using AOPI stain following manufacturer’s recommended conditions. Single-cell encapsulations were generated utilizing 5’ v1 Gem beads from 10x Genomics (Pleasanton, CA, USA) on a 10x Chromium controller and downstream TCR, and Surface marker libraries were made following manufacturer recommended conditions. All libraries were quantified on a BioRad CFX 384 (Hercules, CA, USA) using Kapa Biosystems (Wilmington, MA, USA) library quantified kits and pooled at an equimolar ratio. TCRs, surface markers, and tetramer generated libraries were sequenced on Illumina (San Diego, CA, USA) NextSeq550 instruments. Sequencing data were processed using the Cell Ranger Software Suite (Version 3). Samples were demultiplexed and unique molecular identifier (UMI) counts were quantified for TCRs, tetramers, and gene expression.

**Single-cell Transcriptomic Analysis.** Hydrogel-based RNA-seq data were analyzed using the Cell Ranger package from 10X Genomics (v3.1.0) with the GRCh38 human expression reference (v3.0.0). Except where noted, Scanpy (v1.6.0,(*56*)) was used to perform the subsequent single cell analyses. Any exogenous control cells identified by TCR clonotype were removed before further gene expression processing. Hydrogels that contain UMIs for less than 300 genes were excluded. Genes that were detected in less than 3 cells were also excluded from further analysis. Several additional quality control thresholds were also enforced. To remove data generated from cells likely to be damaged, upper thresholds were set for percent UMIs arising from mitochondrial genes (13%). To exclude data likely arising from multiple cells captured in a single drop, upper thresholds were set for total UMI counts based on individual distributions from each encapsulation (from 1500 to 3000 UMIs). A lower threshold of 10% was set for UMIs arising from ribosomal protein genes. Finally, an upper threshold of 5% of UMIs was set for the MALAT1 gene. Any hydrogel outside of any of the thresholds was omitted from further analysis. A total of 15,683 hydrogels were carried forward. Gene expression data were normalized to counts per 10,000 UMIs per cell (CP10K) followed by log1p transformation: ln(CP10K + 1).

Highly variable genes were identified (1,567) and scaled to have a mean of zero and unit variance. They were then provided to scanorama (v1.7,(*57*)) to perform batch integration and dimension reduction. The data were used to generate the nearest neighbor graph which was in turn used to generate a UMAP representation that was used for Leiden clustering. The hydrogel data (not scaled to mean zero, unit variance, and before extraction of highly variable genes) were labeled with cluster membership and provided to SingleR (v1.4.0, (*58*)) using the following references from Celldex (v1.0.0, (*58*)): Monaco Immune Data, Database Immune Cell Expression Data, and Blueprint Encode Data. SingleR was used to annotate the clusters with their best-fit match from the cell types in the references. Clusters that yielded cell types other than types of the T Cell lineage were removed from consideration and the process was repeated starting from the batch integration step. The best-fit annotations from SingleR after the second round of clustering and the annotation was assigned as putative labels for each Leiden cluster. Further clustering of transcriptomic data was performed across the genes shown in Figure 5 using KMeans in sklearn (v0.24) with n_clusters set to 8. As the method has a preference to assign like-sized clusters, further consolidation of two central memory clusters was performed.

In order to provide corroboration for the SingleR best-fit annotations and further evidence as to the phenotype of the clusters, gene panels representing functional categories (Naïve, Effector, Memory, Exhaustion, Proliferation) were used to score each hydrogel’s expression profiles using scanpy’s “score_genes” function (*56*) which compares the mean expression values of the target gene set against a larger set of randomly chosen genes that represent background expression levels. The gene panels for each class were: Naïve - TCF7, LEF1, CCR7; Effector - GZMB, PRF1, GNLY; Memory - AQP3, CD69, GZMK; Exhaustion - PDCD1, TIGIT, LAG3; Proliferation - MKI67, TYMS. The gene expression matrix for all hydrogels were first imputed using the MAGIC algorithm (v2.0.4, (*59*)). These functional scores were the only data generated from imputed expression values.

**Scoring pMHC-TCR interactions.** Tetramer data analysis was performed using Python (v3.7.3). For each single-cell encapsulation, tetramer UMI counts (columns) were matrixed by cell (rows) and log-transformed. Duplicates of this matrix were independently Z-score transformed by row or column, and subsequently median-centered by the opposite axis (column or row), respectively. For each pMHC-cell interaction, this provided two scores - inter-tetramer ($S_{tet}$) and inter-cell ($S_{cell}$), which were used to calculate a classifier for unique CDR3 α/β clonotypes across $N$ cells as ${N*\bar{S}}_{tet}*\bar{S}_{cell}$. A classifier threshold of 40 for positive interactions.

**TCR Network Analysis.** TCR motif analysis was performed using scirpy (v0.6.1) with receptor_arms = “any,” metric = “alignment,” and default cutoff of 10. Once clusters were identified, sequence alignment was performed using the pairwise2 module in Biopython (v1.78) and visualized using logomaker (v0.8).

**Recombinant TCR validation.** Recombinant TCRs identified from patient samples were ordered from TWIST Biosciences in the pLVX-EF1a lentiviral backbone (Takara) as a bicistronic TCRβ-T2A-TCRα vector. Viral supernatants from transfected HEK 293T cells were collected 48 and 72 hours after transfection and added to the parental TCRαβ^-/-^ Jurkat J76 cell like (*33*) expressing CD8 and an NFAT-GFP reporter, referred to as J76-CD8-NFAT-GFP. Recombinant TCR surface expression was confirmed through flow cytometry by staining transduced J76-CD8-NFAT-GFP cells with anti-CD3-PE (Clone UCHT1) and anti-TCRab-APC (Clone IP26) antibodies.

To assess functional activity of recombinant TCRs, J76-CD8-NFAT-GFP expressing recombinant TCRs were incubated at a 1:1 ratio with the HLA-A*02:01^+^and HLA-B*07:02^+^ HCC 1428 BL (ATCC CRL-2327) lymphoblastic cell line, with a final concentration of 0.5% DMSO (vehicle) or 50 mM of cognate peptide (New England Peptide, >95% pure). Cell mixtures were incubated in the Sartorius IncuCyte at 37°C, 5% CO_2_ overnight and analyzed for NFAT-GFP expression measured as total integrated intensity (GCU x μm^2^/image) at 12 hours after assay setup. At 16 hours, cells were removed from the IncuCyte and subsequently washed and blocked with BD Staining Buffer (BD 554656), stained with anti-CD3-PE-Cy7 (Clone UCHT1) and anti-CD69-APC (Clone FN50) antibodies, and analyzed using the Intellicyt iQue Screener Plus and FlowJo v10. CD69 activity was measured as percent positive of CD3^+^ cells.

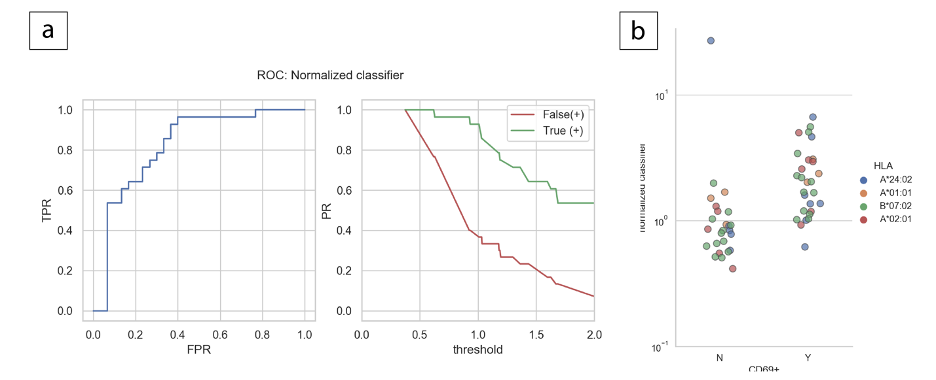

Fig. S1.

Receiver-operator analysis for TCR-pHLA hit identification, where true positives were determined by CD69+ readout in recombinant TCR assays. (a) Receiver-operator curve showing true positive rate (TPR) versus false positive rate (FPR), and associated TPR (green) and FPR (red) versus classifier threshold. (b) Shown are distributions of classifiers for interactions found to be negative (N) by CD69 assay or positive (Y).

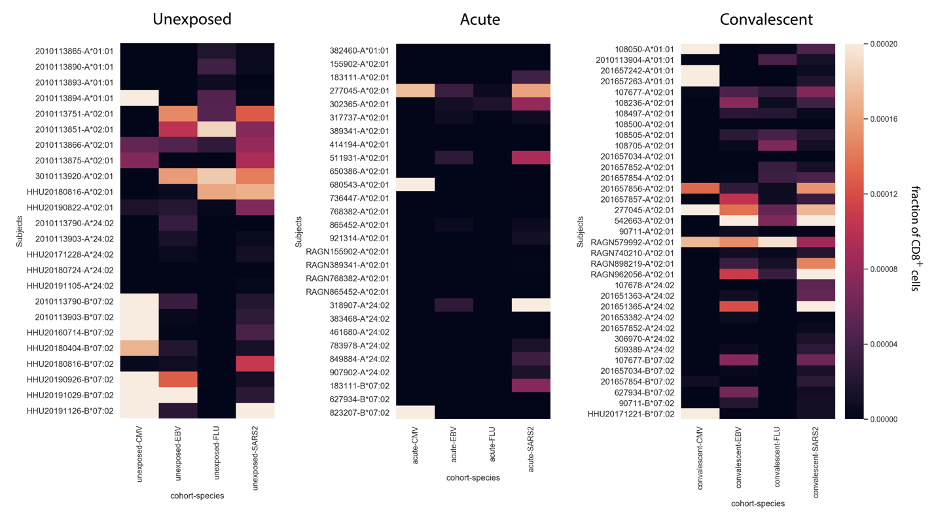

Fig. S2.

Overall reactivity to CMV, EBV, influenza, and SARS-CoV-2 by subject by cohort. Shown are cumulative fractions of T cells found to be reactive to any epitope assayed within the designated species.

Table S4.

Putative hits identified by antigen, represented the number of clonotypes that were found to be specific to any epitope within the given antigen.

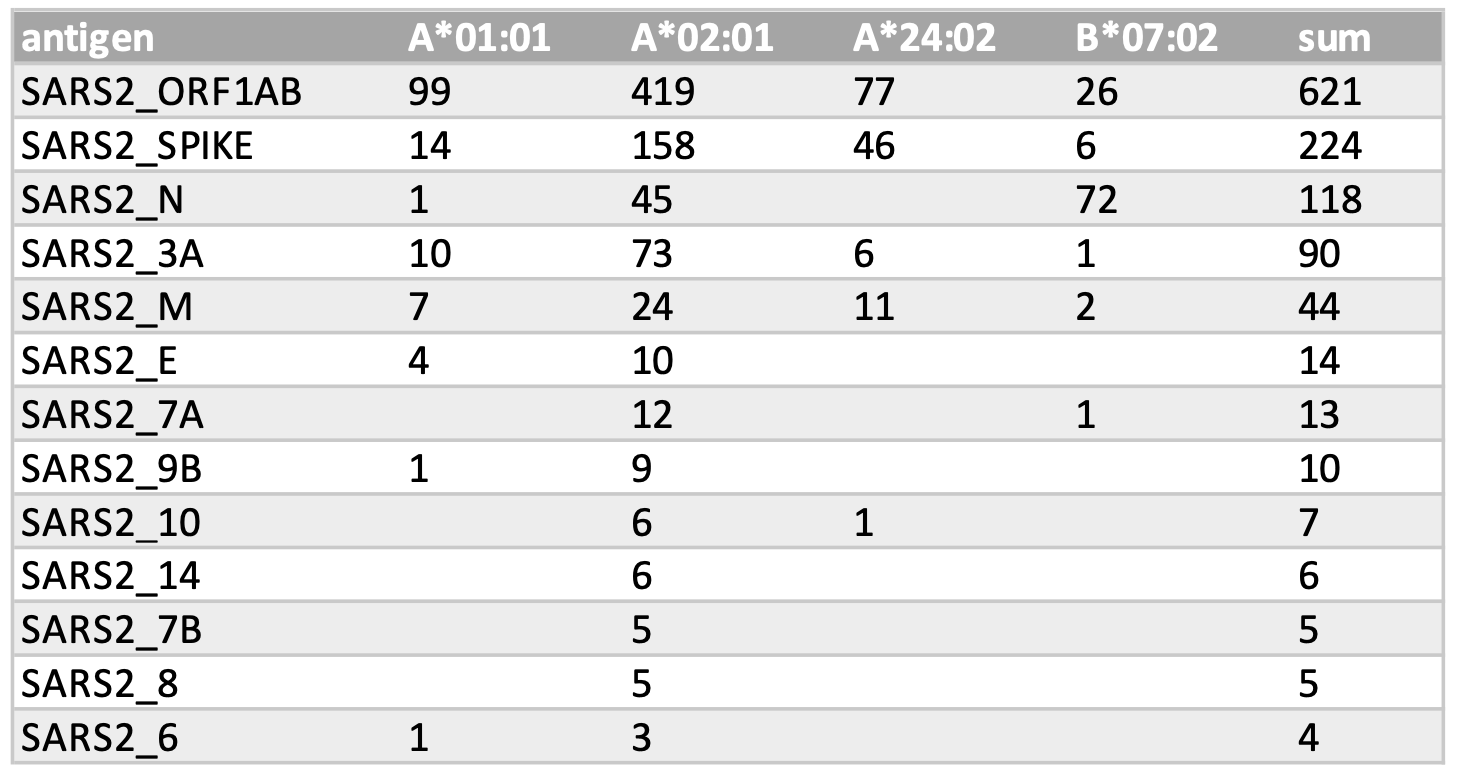

Table S5.

Epitopes and hits discovered in this work that would be impacted by high-threat SARS-CoV-2 variants. Shown are the epitopes included in this study that align with the specified mutation along with the number of clonotypes found to bind that epitope across the specified number of cells.

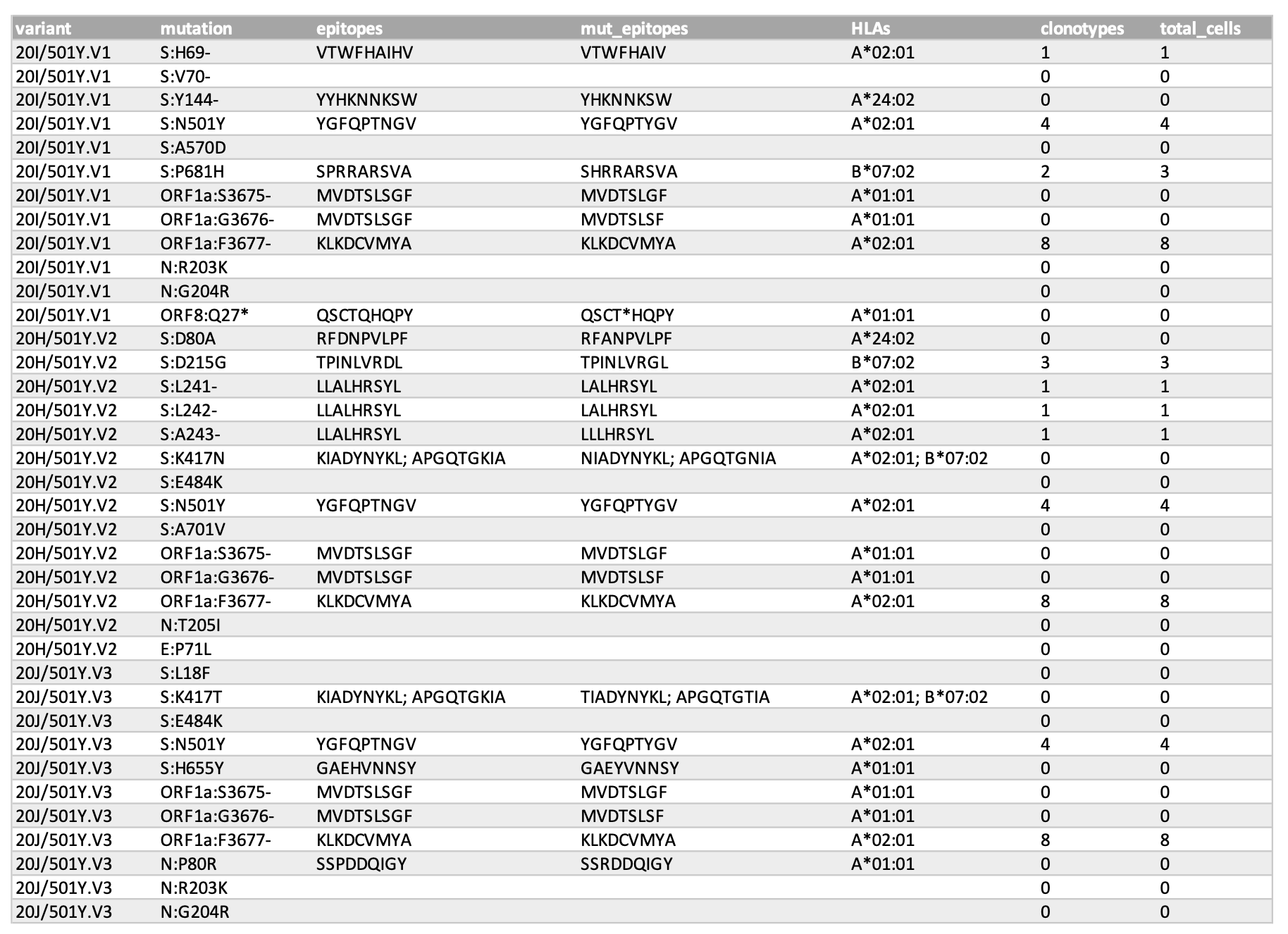

Table S6.

Recombinant TCR-peptide-HLA interactions confirmed by functional testing. Shown are the HLA, epitope, and alpha/beta CDR3 sequences for interactions originally identified across nCells.

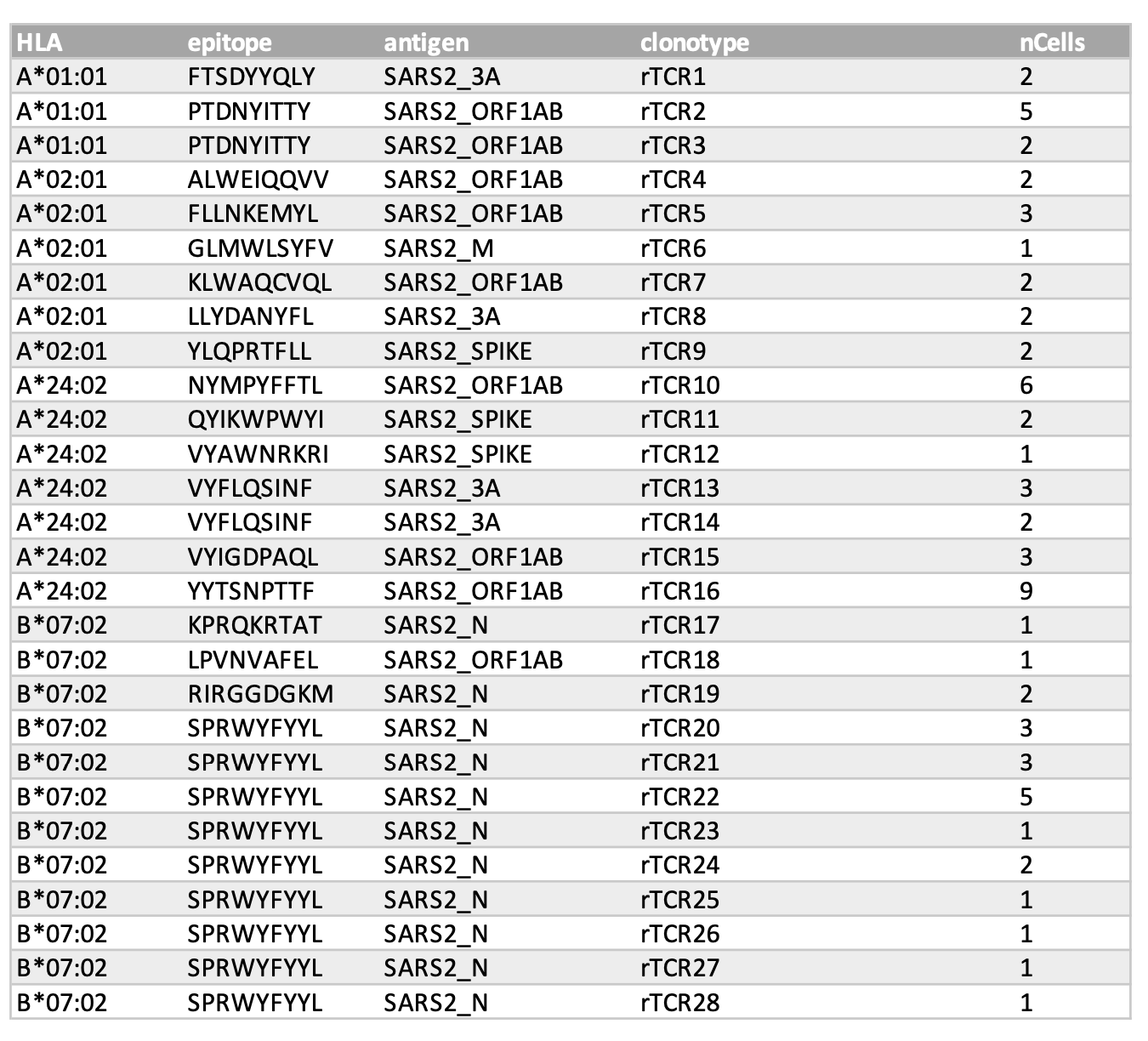

Table S1.

Subject summary and metadata.

Table S2.

Putative TCR-peptide-HLA hits identified in this study.

Table S3.

Summary of SARS-CoV-2 epitopes identified in this study compared to those obtained in other publications.

Table S7.

Representation of transcriptional phenotype by epitope specificity and cohort. These values are consistent with the cells/clusters represented in Figure 5 of the main text.
